# Hyphosphere Hunting Grounds: Fungal Hyphae Guide a Protist Predator to its Bacterial Prey

**DOI:** 10.64898/2026.11.22.768452

**Authors:** Gayan Abeysinghe, Esau De la Vega Camarillo, Shravan Sharma Parunandi, Stephen Taerum, Muthukumar Bagavathiannan, Lindsay Triplett, Sanjay Antony-Babu

## Abstract

Interactions between microeukaryotes mediated by prokaryotic intermediaries remain poorly understood. Here, we show that the hyphae of a plant pathogen guide bacterivorous protists into bacterial habitats, transforming a fungal dispersal route into a potential hunting ground. Although fungal hyphae are established dispersal corridors for bacteria, their capacity to guide individual protist cells remains largely unexplored. Time-lapse imaging of *Fusarium oxysporum* f. sp. *vasinfectum* Race 4 revealed bacteria and the flagellate *Paracercomonas* sp. moving bidirectionally along hyphae with significant axial alignment, despite contrasting motility regimes: rapid, superdiffusive bacterial movement and slower, subdiffusive protist movement. The ciliate *Paracolpoda steinii* also exhibited directional alignment along an inert glass rod, demonstrating that physical geometry can bias protist movement independently of fungal-specific cues. Across successive generational culturing of hyphal-baits, protist communities converged reproducibly toward *Paracolpoda* dominance, whereas bacterial assembly was largely stochastic, revealing contrasting assembly processes within the same habitat. In a dual-microcosm assay, fungal bridges enabled bacteria and protists to cross a 3 mm air gap, while the presence of *P. steinii* reduced bacterial arrival approximately 8-fold, consistent with predator-associated filtering. Together, these findings extend the fungal-highway concept beyond microbial transport, identifying pathogenic fungal hyphae as multi-kingdom ecological corridors where physical guidance, community assembly and predator-associated effects on dispersal intersect.

## Introduction

Fungal hyphae connect soil pore space that is otherwise discontinuous. Especially, through their sustained apical growth, fungal mycelia form networks that bridge spatially separated soil microhabitats over centimeter and potentially larger scales. Thin liquid films associated with hyphal surfaces further increase hydraulic connectivity, allowing water-dependent microorganisms to traverse otherwise disconnected pore spaces (Ritz & Young, 2004; Allen, 2007; Dechesne et al., 2010). Direct physical association with hyphae is also a widespread mode of bacterial-fungal interaction across environmental, agricultural, and host-associated systems (Deveau et al., 2018; Steffan et al., 2020). In laboratory systems, motile bacteria have dispersed along fungal hyphae at rates approaching 1 cm per day, a dispersal mechanism subsequently termed the “fungal highway” (Kohlmeier et al., 2005) and since documented across diverse fungal and bacterial taxa and soil environments (Warmink et al., 2011; Simon et al., 2015; Simon et al., 2017), with experimental platforms subsequently developed to recruit and isolate hyphae-associated bacteria directly from environmental samples (Junier et al., 2021; Kelliher et al., 2025). A well-characterized example further demonstrated that the interaction can be reciprocal where *Bacillus subtilis* dispersed along *Aspergillus nidulans* mycelia while supplying thiamine that promoted fungal growth, making the highway a mutualistic exchange rather than simple bacterial exploitation of fungal infrastructure (Abeysinghe et al., 2020).

This ecology has been characterized largely in saprotrophic and mycorrhizal fungi, whose hyphae facilitate prokaryotic dispersal through spatially fragmented soil habitats and create selective niches for associated microorganisms (Jansa et al., 2013; Jansa & Hodge, 2021; Kuhn et al., 2022; Krug Vieira et al., 2025). The hyphosphere is a well-recognized interaction habitat that recruits bacteria to its vicinity, where they exchange nutrients with their fungal scaffold and establish niches around fungal surfaces, though it remains far less characterized than the rhizosphere (Frey-Klett et al., 2011; de Boer et al., 2005; Wang et al., 2024). However, the hyphosphere of a plant pathogen poses a different question. Microorganisms travelling along pathogenic hyphae form part of a disease-relevant microbial assemblage, placing fungal-highway dispersal within a pathobiome context in which interactions among the pathogen, associated microorganisms, host, and environment may influence disease expression (Vayssier-Taussat et al., 2014; Bass et al., 2019). This spatially resolved interface has recently been conceptualized as the pathomycosphere, the hypha-associated microbial assemblage of a fungal pathogen where direct interactions with associated microorganisms can influence pathogen biology and disease ecology (Abeysinghe and Antony-Babu, 2026). We previously identified distinct and recurrently associated bacterial communities in the hyphospheres of *Fusarium oxysporum* f. sp. *niveum* and f. sp. *cubense* (Thomas & Antony-Babu, 2024; Thomas et al., 2025). We also showed that *F. oxysporum* f. sp. *vasinfectum* Race 4 (FOV4), a highly virulent cotton wilt pathotype first reported causing disease in Texas independently of prior recognized nematode wounding (Halpern et al., 2018; Wagner et al., 2022), recruits a *Pseudomonas*-dominated hyphosphere containing isolates that promote FOV4 hyphal extension (Antony-Babu et al., 2025). Methods used in these previous studies did not allow examining organisms beyond bacteria.

In the study presented here, we show soil protists as additional participants. As major consumers of soil bacteria, protists exert top-down control on bacterial communities and drive nutrient turnover through the microbial loop (Bonkowski, 2004; Geisen et al., 2018; Gao et al., 2019; Martins et al., 2022), and some are also mycophagous, consuming fungal hyphae, spores, or yeasts directly (Geisen et al., 2016). This places a protist in functional loggerheads with the hyphosphere itself, which supports bacterial growth and recruits bacteria to the hyphal surface, since a bacterivore arriving there removes what the hypha has gathered. In AMF systems, the nutrient environment shapes both prokaryotic and eukaryotic community assembly in the hyphosphere (Faghihinia et al., 2024). In a related experimental system, adding the protist *Polysphondylium pallidum* to a chitin-degrading bacterium increased organic-nitrogen transfer through the fungus to the host plant by more than 60% relative to the bacterium-only condition, through stimulation of the microbial loop (Rozmoš et al., 2022). Ciliates of the order Colpodida can also redistribute beneficial bacteria along growing plant root systems (Hawxhurst et al., 2023). Protists are therefore plausible active components of the hyphosphere, with the capacity to alter bacterial distribution, trophic interactions, and nutrient turnover.

Whether individual protist cells are guided along the hyphal surface, however, has never been directly demonstrated. Hyphae increase protist abundance and dispersal range across otherwise disconnected soil pores (Mafla-Endara et al., 2021), and specific protist taxa are enriched in association with AMF hyphae (Zhang et al., 2024). These findings establish the protist-hyphal interactions biology at a population and compositional level, and opens the question of what individual cells do on that surface: whether a protist uses the hypha directly as a locomotory guide, or merely benefits from hypha-induced hydration and altered pore connectivity. The uncertainty is also biophysical. Especially because protists differ markedly from bacteria in cell size and locomotory machinery, bacterial fungal-highway behavior cannot be assumed to extend to protist cells.

Here we show that individual protists can use fungal hyphae as locomotory surfaces, and that directional guidance can also emerge on an inert structure of comparable geometry. Time-lapse imaging of FOV4 hyphae recovered from hyphal-baiting microcosms showed both bacteria and a flagellate protist provisionally assigned to *Paracercomonas* sp. moving with significant alignment to the hyphal axis and net displacement along it, bidirectionally, with bacteria moving 4.4-fold faster than the protist. To test whether this guidance requires the living fungus, we tracked a second, ciliate protist (*Paracolpoda steinii*) on both FOV4 hyphae and an inert glass rod used as a nonliving linear surface control. On the rod, movement was significantly more aligned with its long axis than expected by chance; on real hyphae, alignment with local branch geometry did not differ from chance, a pattern that may reflect more exploratory movement on a branched, biologically active surface supporting an associated microbial community rather than passive alignment to a uniform rod. Nanopore metabarcoding across successive hyphal-baiting generations showed convergent assembly of both bacterial and protist communities, but by different routes: replicate protist community richness reproducibly reduced to a single ciliate species, while bacterial assembly was dominated by stochastic, dispersal-mediated homogenization. In a dual-microcosm assay, a fungal bridge was required for either bacteria or protists to colonize a recipient microhabitat across a 3 mm air gap. Bacterial colonization was reduced approximately eightfold when the protist was present, consistent with, although not demonstrating, grazing-mediated filtering of the bacteria that completed the crossing. These results indicate that protists are not incidental passengers on fungal highways but active participants that can exploit the same infrastructure their bacterial prey use to track and filter them, a mode of microeukaryote interaction distinct from simple habitat association or direct antagonistic contact.

## Materials and Methods

### Hyphal-recruitment microcosm and generational culturing

As described in Abeysinghe et al., (2026) soil was collected from an actively cultivated cotton field at the Texas A&M AgriLife Research Field, Brazos Bottom, Texas, USA (30°32′14.0″ N, 96°25′40.1″ W), characterized by clay loam soil. Surface litter and plant residues were removed prior to sampling. Soil was collected from a depth of 5-15 cm at 10 randomly selected locations, combined in sterile bags, homogenized, and transported to the laboratory on ice within 2 h. To prepare the soil-tea inoculum, 6 g of homogenized soil was suspended in 10 mL of sterile 1× phosphate-buffered saline (PBS) supplemented with 30% (v/v) glycerol and vortexed for 5 min at room temperature to detach soil-associated microorganisms. The resulting suspension was used as the microbial inoculum for all microcosms.

Pima cotton seeds (*Gossypium barbadense* cv. Pima S7) were surface sterilized by immersion in 1.25% (w/v) NaOCl for 3 min, followed by 70% (v/v) ethanol for 3 min, rinsed three times with sterile reverse-osmosis water, and pre-germinated on 2% (w/v) Gelrite in the dark at room temperature for 48 h.

Microcosms were prepared using a 3:1 mixture of autoclaved field soil and sterile sand. Field soil was sterilized at 121 °C for 60 min on three consecutive days before mixing with sterile sand. The soil mixture was dispensed into sterile 450-mL plastic pots and inoculated with 5 mL of soil-tea suspension. A single pre-germinated cotton seedling was aseptically planted in each pot. Five independent biological replicate microcosms were established.

Hyphal-baiting assemblies consisted of sterile 100-µm cell strainers (Falcon CLS352360, Corning, USA) fitted with a sterile glass coverslip at the base to minimize direct soil entry. A 1.5x 1.5 cm agar plug excised from the actively growing margin of a 3-day-old culture of *Fusarium oxysporum* f. sp. *vasinfectum* race 4 isolate F4AB grown on 0.1x potato dextrose agar was placed inside the cell strainer with the colony margin positioned approximately 1 mm from the inner wall. The assembly was inserted approximately 3 cm into the soil and positioned approximately 2 cm from the seedling base, with the fungal colony facing the hypocotyl, allowing hyphae to extend through the mesh into the surrounding rhizosphere while maintaining a recoverable boundary between the fungal inoculum and the soil environment. Each microcosm received approximately 50 mL of sterile 0.25x Hoagland’s solution and was maintained in a growth chamber under a 12.5 h light/11.5 h dark photoperiod at 23 °C (day) and 18 °C (night).

One week after introduction of the hyphal-baiting assembly, the cell strainer was removed aseptically. Soil particles adhering to the outer strainer wall and mesh and bound by hyphae extending into the surrounding rhizosphere were gently collected using sterile forceps and placed onto 0.1x potato dextrose agar (1.5% w/v agar). Plates were incubated at 28 °C until actively growing hyphae emerged from the collected hyphal-associated soil aggregates. Hyphal tips were then aseptically excised and introduced into fresh hyphal-recruitment microcosms following the same procedure described above to generate the first generation (G1). The same recruitment, recovery, and hyphal-tip transfer procedure was repeated to generate the second generation (G2). Rhizosphere samples from the initial planted microcosms and hyphal-tip-associated samples from G1 and G2 were collected from five independent biological replicates at each stage for DNA extraction, PCR amplification, and full-length 16S and 18S rRNA gene metabarcoding.

### DNA extraction, amplicon sequencing, and taxonomic classification

Hyphal-tip samples (∼5 mm segments) were excised aseptically, flash-frozen in liquid nitrogen, and stored at -80 °C; rhizosphere soil particles (0.5 g) were frozen similarly. Total genomic DNA was extracted using the ZymoBIOMICS DNA Miniprep Kit (Zymo Research). The near-full-length 16S rRNA gene (∼1,500 bp; primers 27F/1492R) and 18S rRNA gene (∼1,800 bp; primers F1/R1) were amplified with KAPA2G Fast HotStart ReadyMix, purified with AMPure XP beads, and normalized to equal DNA mass prior to sequencing on an Oxford Nanopore PromethION platform (R10.4.1 chemistry, Native Barcoding Kit 24 V14) by Plasmidsaurus (Eugene, OR). Reads were basecalled with Dorado (Super-Accurate model), quality-filtered (Q ≥ 10, 400-3,000 bp), and adapter-trimmed with Porechop. Taxonomic classification used EMU against rrnDB + NCBI Targeted Loci (16S) or NCBI Targeted Loci + PR2 (18S). Taxa with mean relative abundance < 0.1%, and reads classified as mitochondria, chloroplasts, Metazoa, or Embryophyta, were excluded; residual fungal 16S assignments (*Balansia*, *Claviceps*, *Beauveria*) and non-target plant/algal 18S genera were removed as non-bacterial or non-protist contaminants.

Full-length 18S rRNA reads classified as *Colpodida* (NCBI taxid 49142) and *Paracercomonas* (taxid 45338) by Emu (Curry et al., 2022) were pooled across all samples and sequencing runs after removing reads assigned to Bacteria. Because a single Nanopore read can carry errors or chimeric artifacts, we did not use any individual read as the representative sequence for each taxon. Instead, for each taxon we retained only reads classified with high certainty whose length fell within ±5% of the median read length for that taxon, ranked them by Emu confidence score, and selected the top 40 reads. These reads were aligned with MAFFT v.7 (Katoh and Standley, 2013), and a consensus sequence was built from the alignment by majority rule at each column. Alignment columns in which more than half of the reads had a gap were excluded, so that insertions present in only a minority of reads were not carried into the consensus. Consensus calling was done with a custom Python script using Biopython v1.88 (Cock et al., 2009). Since chimeric reads are single-molecule events that are unlikely to recur across dozens of independent reads, this approach reduces the risk that a chimera or sequencing error ends up in the final sequence. The resulting consensus sequences were used as the query sequences for the phylogenetic analysis described below.

Two sequence datasets were generated: one contained the putative *Paracolpoda* consensus sequence and 42 reference sequences from the Protist Ribosomal Reference (PR^2^) database (37 from the order Colpodida, five from the order Cyrtolophosidida as an outgroup), while the other contained the *Paracercomonas* consensus sequence and 23 reference sequences from PR^2^ (19 from the family Paracercomonadidae, four from the family Cercomonadidae as an outgroup). Each dataset was aligned using the Auto strategy in MAFFT v.7 (Katoh and Standley 2013). The datasets were then visualized for regions of ambiguous alignments, and trimmed as necessary. The final Colpodida and Paracercomonadidae alignments were 1684 and 1704 bp, respectively. Maximum likelihood (ML) analyses were conducted using RaxML v.8.2.10 (Stamatakis 2014) following a GTR+Γ model, and branch support was generated using 1000 bootstrap replicates. Bayesian inference (BI) analyses were conducted using MrBayes v.3.2.7 (Ronquist et al. 2012) following the same evolutionary model used for ML analyses. BI analyses were conducted with four Markov chain Monte Carlo chains up to 5,000,000 generations. Tracer v.1.7.2 (Rambaut et al. 2018) was used to determine burn-in, and trees sampled in the burn-in phase were discarded. Trees were sampled every 100 generations after which posterior probabilities were calculated.

### Microscopy, motility tracking and kinematics

#### Hyphal motility assay

Bacterial and protist motility on hyphae was imaged directly on the 0.1x PDA plates from the G1 and G2 hyphal-baiting generations (see hyphal-recruitment microcosm and generational culturing). On these plates, actively growing FOV4 hyphae emerged together with the co-recovered bacteria and protists. Hyphae were imaged in situ once they had extended from the aggregates; the plates received no further manipulation.

#### Glass-rod surface assay

A glass rod (1 mm diameter x 10 cm; World Precision Instruments, Sarasota, FL, USA) was used as a non-living linear surface control. Its diameter is approximately two orders of magnitude wider than an FOV4 hypha. *P. steinii* was isolated together with its associated bacteria from the G2 recovery plates. Antibiotic treatments failed to free the protist of bacteria, so it was maintained and used as a protist-bacteria co-culture. The co-culture was grown in tryptic soy broth (TSB) at 28 °C for 48 h. Cells were then washed with sterile 1x PBS followed by sterile water and resuspended in sterile water to an OD₆₀₀ of 0.1. Glass rods were surface-sterilized with 10% sodium hypochlorite followed by 70% (v/v) ethanol, rinsed with sterile water and air dried. The co-culture suspension (1 µL) was spotted at the starting point on 0.1x PDA Petri plate, and the rod was placed with one end in contact with the inoculum. Plates were kept closed and incubated at 28 °C for 48 h, after which protist movement was imaged along the rod surface. Five independent replicate plates were prepared.

#### Image acquisition

Bacterial, Protists movement along FOV4 hyphae, and *P. steinii* movement on FOV4 hyphae versus the glass rod, were recorded using an Echo Revolution microscope (Echo, San Diego, CA, USA) in upright configuration. Bright-field images were acquired with a 20×0.45 NA Plan Fluorite objective (OFN22). Plates were imaged with lids closed at room temperature. All the images and time-lapse sequences were acquired using the native Echo acquisition software. Only linear adjustments to brightness and contrast were applied, uniformly across all samples. Scale bars were generated automatically from the calibrated optical magnification.

#### Trajectory analysis

Cell trajectories were generated using the TrackMate plugin in FIJI/ImageJ. For each track, speed and net displacement were calculated from calibrated frame to frame centroid positions. Track straightness was quantified as the directionality ratio (DR), the ratio of net end to end displacement to total path length, ranging from 0 (purely diffusive) to 1 (perfectly straight). Mean squared displacement (MSD) was calculated per track and fit to a power law (MSD) to classify motion as sub-diffusive (α < 1), Brownian (α = 1) or super-diffusive (α > 1).

Angular deviation was calculated as the angle between each track’s net displacement vector and the local guiding axis, folded to 0-90°, so that tip-ward and base-ward movement were treated equivalently. Alignment was tested against random expectation in two ways, according to surface geometry. (i) For bacteria and *Paracercomonas* on single, continuously traceable hyphae, deviation was measured from the hyphal long axis. Isotropic random movement gives an expected mean deviation of 45° for a single axis, so the observed mean was compared with 45° using a one-sample t-test. Results were additionally confirmed with an exact one-sample Wilcoxon signed-rank test, which makes no assumption of normality. (ii) For *P. steinii*, deviation was measured from the rod’s single long axis, or on branched hyphae from the nearest of three manually traced local branches. Choosing the nearest of several axes lowers the deviation expected by chance below 45°, and no analytical expectation exists for this geometry. Each condition was therefore tested against a simulated random baseline (rod, 45.0°; branched hyphae, 17.2°) using a track-length-matched permutation test (200,000 iterations per condition, matching the number and lengths of observed tracks). Two-tailed p-values were calculated as the proportion of simulated means at least as extreme as the observed mean.

All comparisons between two groups (bacteria vs. *Paracercomonas*; glass rod vs. hyphae) of speed, net displacement, DR and angular deviation used two-tailed Mann-Whitney U tests.

#### Scanning electron microscopy

The cultures grown on agar medium was fixed in Trump’s fixative (McDowell and Trump, 1976) for 24 h at room temperature. All subsequent washing and dehydration steps were performed using Pelco Biowave microwave processor (Ted Pella, Redding, CA, USA), at 150W power setting and the ColdSpot plate temperature set to 20 °C. In each step, the sample was microwaved for 1 minute, then allowed to rest for 5 minutes and microwaved for 1 minute again. All washing steps were performed under vacuum, while dehydration and hexamathyldisilazane infiltration was done at normal atmospheric pressure (Ferris et al., 2009). The fixed sample was washed three times in Trump’s buffer, post-fixed in 1% aqueous osmium tetroxide for 1 hour at room temperature, washed in water three times and dehydrated in ethanol series (30,50, 70, 95, 95% v/v). Afterwards, dehydration in 100% ethanol was repeated three times with the resting period extended to 1-2 hours. Samples were then gradually transitioned to hexamethyldisilazane (HMDS) drying agent (Bhattacharya et al., 2020): Ethanol:DMDS 2:1, 1+2, then HMDS alone three times, with microwave treatment as above, resting period 45 minutes in each step. The sample was then allowed to air dry overnight, then dried under vacuum at room temperature for additional 2 days. The dry sample was removed from the Petri dish, broken into smaller pieces, mounted on 25 mm SEM pin mounts and sputter coated with 10 nm Platinum/Palladium (80/20 Pt/Pd mix). Imaging was performed using FEI Quanta 600 FESEM and 10 kV Accelerating voltage. Sample preparation and imaging was performed by the Microscopy and Imaging Center Core Facility at Texas A&M University (RRID:SCR_022128).

#### Dual-microcosm dispersal assay

To test whether fungal hyphae are required for the cross-kingdom dispersal of bacteria and protists, we developed a dual-microcosm assay in which a donor and a recipient compartment are separated by an air gap (Figure 4a). The gap prevents transfer by diffusion or swimming through a continuous liquid film. Colonization of the recipient compartment is therefore possible only if hyphae grow across the gap and form a bridge.

Each microcosm pair consisted of two sterile 100 µm cell strainers (Falcon, Corning, USA) placed side by side with a 3 mm air gap between their outer mesh walls. As in the hyphal-baiting assemblies, a sterile round glass coverslip was fitted at the base of each strainer so that hyphae could not grow underneath. A 1 cm diameter disc of donor culture on 0.1x PDA was placed inside the donor strainer, at the edge facing the gap. A sterile 1 cm diameter 0.1x PDA disc was placed at the facing edge of the recipient strainer. Hyphae growing from the donor disc could therefore reach the recipient disc only by passing through both mesh walls and across the air gap.

Donor discs were prepared for four treatments: (i) FOV4 + protist + bacteria, (ii) FOV4 + bacteria, (iii) protist + bacteria without FOV4, and (iv) bacteria only. An FOV4 only control was also included. Plates were inoculated with FOV4 (1 µL of a 10³ conidia mL⁻¹ suspension), the bacterial suspension (OD₆₀₀ 0.1), and the protist-bacteria coculture (OD₆₀₀ 0.1), according to treatment. Because the protist could not be cultured free of its associated bacteria, a protist-only treatment was not possible.

After 72 h of incubation at 28 °C, the gap was inspected for hyphal bridging. The recipient disc was then examined by bright-field microscopy (Echo Revolution; see Image acquisition) at the point where hyphae entered it. Bacterial cells, protist cells on the recipient disc were counted, and treatments with no cells detected were recorded as not detected (ND). Four independent microcosm pairs were established per treatment. Recipient colonization was compared between treatments by exact two-tailed Mann-Whitney U tests.

#### Community composition and assembly analysis

Genus-level relative abundances were averaged across replicates per generation and visualized as alluvial diagrams of the top genera (18S: 7; 16S: 8, by maximum mean abundance). Alpha diversity (observed richness, Shannon, Simpson, Pielou) was compared across rhizosphere, G1, and G2 by Kruskal-Wallis with FDR-corrected Dunn’s post-hoc tests; beta diversity was assessed by PCoA of Bray-Curtis dissimilarity, tested by PERMANOVA (999 permutations). Community assembly was inferred from the beta nearest-taxon index (β-NTI; Stegen et al., 2013), calculated from MAFFT/IQ-TREE phylogenies with a 999-permutation null model. Comparisons with β-NTI < −2 were classified as homogeneous selection and those with β-NTI > +2 as variable selection. Pairwise comparisons with |β-NTI| ≤ 2, indicating that selection alone could not explain the observed turnover, were further partitioned using the Raup-Crick metric standardized to Bray-Curtis dissimilarities (Rcbray; Stegen et al., 2015). For each such pair, null communities were drawn without replacement from a regional species pool defined across all samples, with taxon selection probability weighted by regional occurrence frequency and community size constrained to match the observed richness of each sample; Bray-Curtis dissimilarity was calculated between each null pair and compared to the observed value across 999 iterations to yield Rcbray, scaled to the interval [-1, 1]. Pairs with Rcbray < -0.95 were classified as homogenizing dispersal, Rcbray > +0.95 as dispersal limited, and Rcbray ≤ 0.95 as undominated (drift).

#### Statistical computing and data availability

All statistical analyses were performed in Python v3.14.4 using SciPy v1.18.1, NumPy v2.5.3, pandas v3.0.6, scikit-bio v0.7.4 and scikit-posthocs v0.17.0. Image processing used OpenCV v5.0.0, and cell tracking used FIJI/ImageJ with the TrackMate plugin v.8.1. All tests were two-tailed, with significance assessed at α = 0.05.

For community data, alpha diversity indices were compared among rhizosphere, G1 and G2 samples by Kruskal-Wallis tests followed by Dunn’s post-hoc tests, with p-values adjusted for multiple comparisons using the Benjamini-Hochberg false discovery rate procedure. Differences in community composition were tested by PERMANOVA on Bray-Curtis dissimilarities with 999 permutations. β-NTI and Rcbray null models were computed with 999 randomizations each as described above.

Raw sequencing reads, consensus sequences and custom analysis scripts have been deposited in figshare (https://doi.org/10.6084/m9.figshare.32896301)

## RESULTS

### Bacteria and a flagellate protist show surface-guided, bidirectional motility along FOV4 hyphae

Time-lapse microscopy of FOV4 hyphae recovered from the first and second generations (G1 and G2) of hyphal-baiting microcosms resolved active movement by both hyphosphere bacteria and protists directly on the hyphal surface (Figures 1a, 2a; Supplementary videos 1-2). The bacterial isolate was provisionally assigned to *Enterobacter* based on colony morphology and the dominant hyphosphere 16S taxon, while the flagellate protist was the dominant hyphosphere flagellate *Paracercomonas marina*. Bacteria moved substantially faster than the protist (mean track velocity 28.2 vs. 6.4 µm s⁻¹, a 4.4-fold difference; Mann-Whitney U, p = 2 x 10^-^⁵; Figure 1b). Both organisms moved with significant axial alignment to the hyphal long axis relative to the 45° expectation under random orientation (bacteria, 11.9° mean deviation, p < 0.0001; protist, 7.5°, p = 0.0009), with bacteria showing modestly but significantly greater deviation than the protist (Mann-Whitney U, p = 0.007; Figure 1c). Space-time trajectories showed both tip-ward and base-ward movement in each organism, indicating bidirectional rather than strictly outward-directed translocation (Figure 1d).

**Figure 1.**
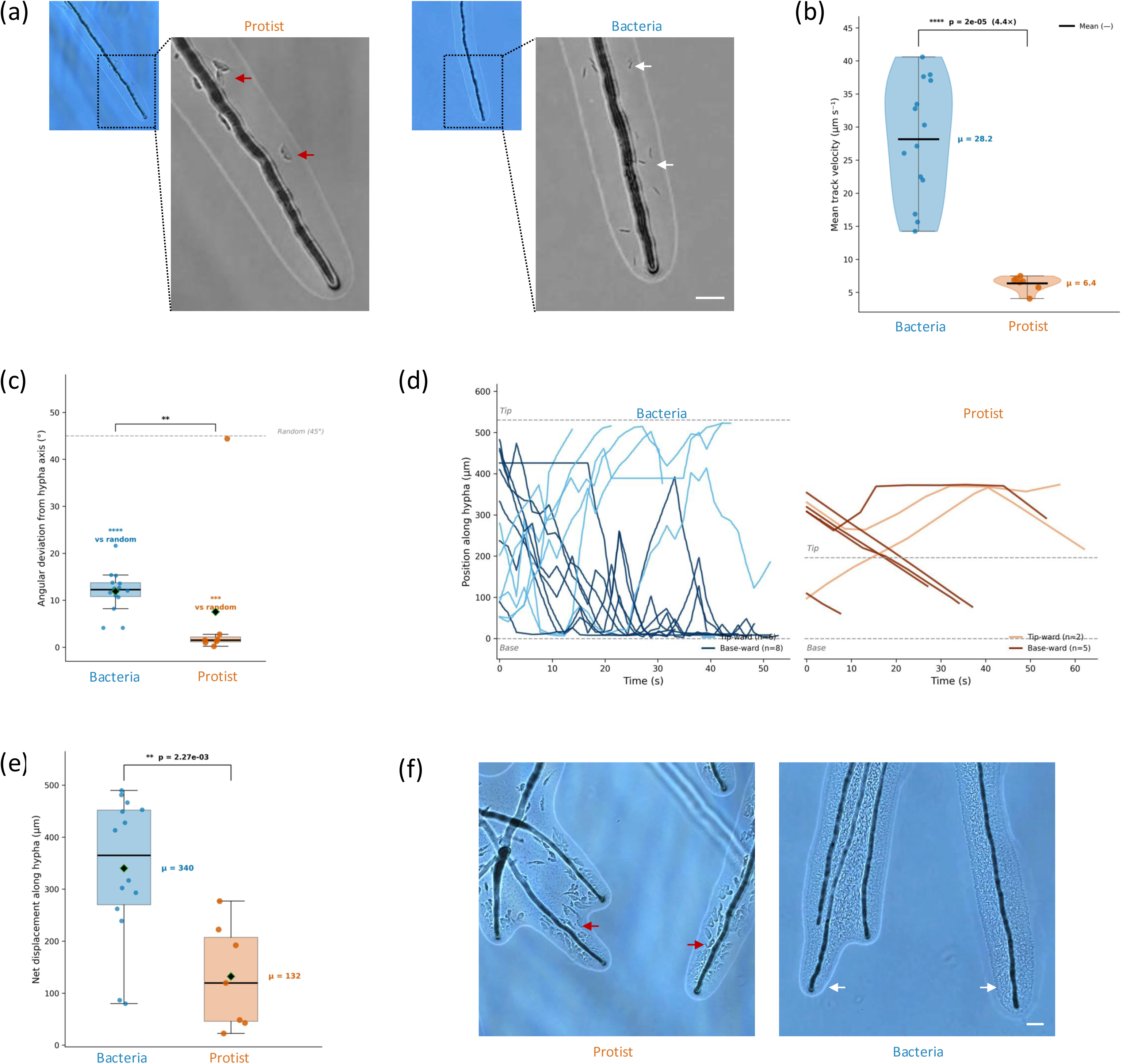
Surface-guided motility of bacteria and flagellate protists along FOV4 fungal hyphae. **(a)** *Fusarium oxysporum* f. sp. *vasinfectum* Race 4 (FOV4) hyphae with bacteria (*Enterobacter* sp., white arrows) and a flagellate protist (*Paracercomonas marina*., red arrows). Scale bar, 10 µm. **(b)** Mean track velocity of bacteria (n = 14) and flagellate protists (n = 7). Horizontal bars, group means. Bacteria moved 4.4-fold faster (28.2 vs. 6.4 µm s⁻¹; Mann-Whitney U, p < 0.0001). **(c)** Angular deviation from the hyphal axis. Dashed line, expected deviation under random movement (45°). Both organisms moved significantly more parallel to the hypha than random (bacteria p < 0.0001, protist p = 0.0009); bacteria showed modestly greater deviation than protists (Mann-Whitney U, p = 0.007). **(d)** Space-time plots of position along the hypha over time, separated by tip-ward (light) and base-ward (dark) movement. Both organisms showed bidirectional motility. **(e)** Net displacement along the hyphal axis. Bacteria showed greater net displacement than protists (340 vs. 132 µm; Mann-Whitney U, p = 0.002). **(f)** Representative micrographs showing accumulation of protists (red arrows) and bacteria (white arrows) at actively extending hyphal tips. Scale bar, 10 µm.

**Figure 2.**
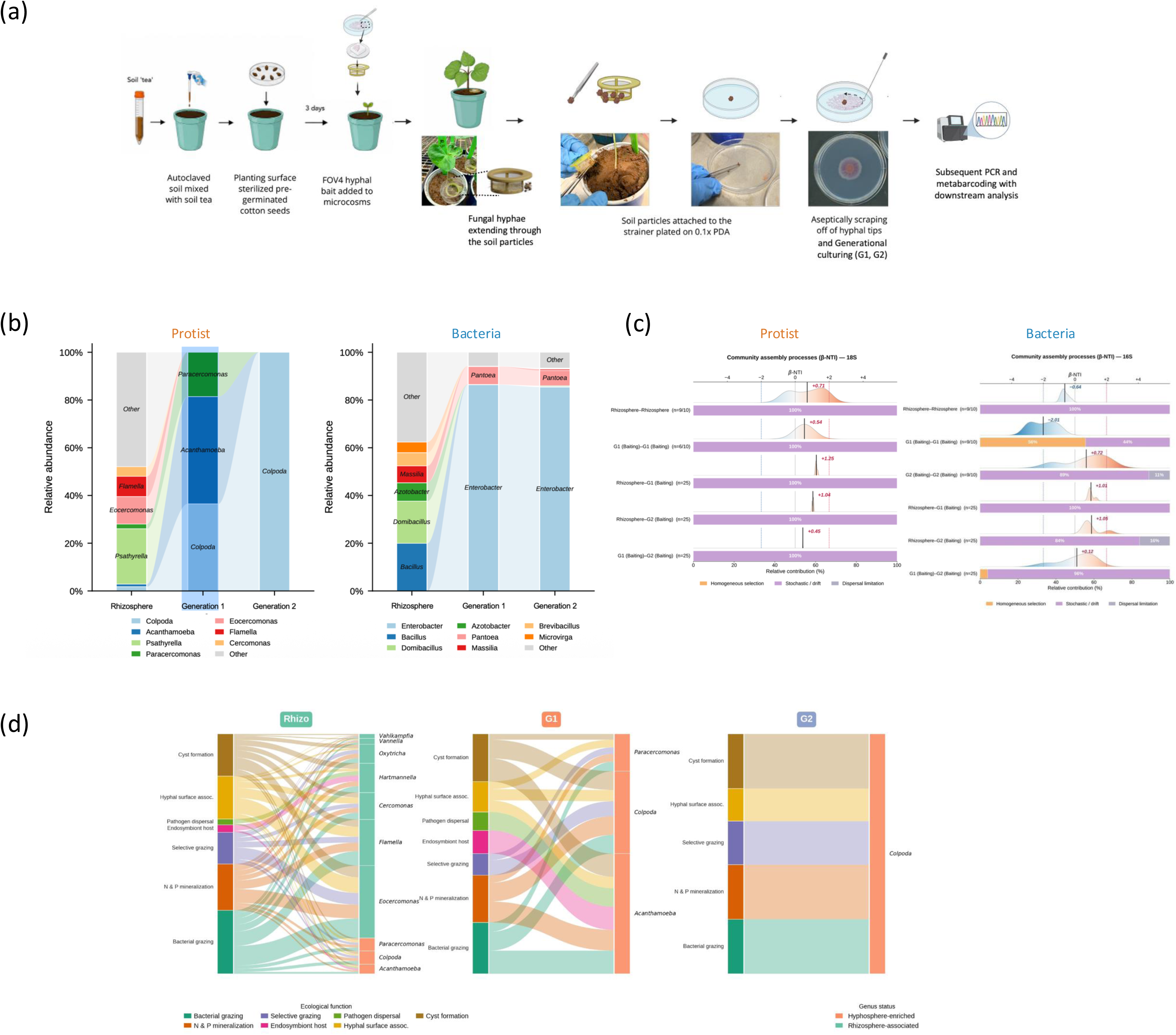
Convergent community succession in the FOV4 hyphosphere across successive hyphal-baiting generations. **(a)** Hyphal-baiting workflow. Autoclaved soil was inoculated with a rhizosphere soil-tea suspension and planted with surface-sterilized, pre-germinated cotton seed. After 3 days, FOV4 hyphal bait was introduced via a sterile cell-strainer microcosm; extending hyphae colonized surrounding soil particles. Hypha-associated particles were collected on the strainer, plated on 0.1× PDA, and hyphal tips were aseptically excised for serial generational culturing (rhizosphere → G1 → G2) prior to DNA extraction, PCR, and metabarcoding. **(b)** Relative abundance of dominant protist (18S, left) and bacterial (16S, right) genera across rhizosphere, G1, and G2. **(c)** Community assembly processes inferred from the β-nearest taxon index (β-NTI), shown as ridge density plots (top) and stacked classification bars (bottom) for protist (left) and bacterial (right) communities, for all pairwise within and between group comparisons (rhizosphere, G1, G2). Pairwise comparisons were classified as homogeneous selection (β-NTI < -2) or variable selection (β-NTI > +2). Comparisons with β-NTI ≤ 2 were further partitioned using RC_bray standardized to Bray-Curtis dissimilarity into homogenizing dispersal (RC_bray < -0.95), dispersal limitation (RC_bray > +0.95), or undominated/drift (RC_bray ≤ 0.95). β-NTI was calculated from MAFFT/IQ-TREE phylogenies with a 999-permutation null model. n = 5 per group; within-group comparisons were drawn from 10 possible pairs and between-group comparisons from 25. Counts shown as n = X/Y indicate pairs excluded because the two communities shared an identical taxon set (β-NTI undefined). **(d)** Ecological function allocation of soil protist genera across rhizosphere (Rhizo), G1, and G2, linking functional category to genus identity and hyphosphere-enrichment status.

Bacteria achieved significantly greater net displacement along the hypha over the imaging window (340 vs. 132 µm; Mann-Whitney U, p = 0.002; Figure 1e) and both organisms were frequently observed at actively extending hyphal tips (Fig. 1f). Kinematic profiling (Supplementary Figure 1) distinguished the character of this motility between kingdoms. Bacterial trajectories were markedly heterogeneous in speed over time, with frequent bursts approaching 100 µm s⁻¹ (Supplementary Figure 1c) and mean squared displacement (MSD) scaled superdiffusively (MSD ∝ τ1.58; Supplementary Figure 1d), consistent with persistent, directed runs interspersed with changes in speed. The protist, by contrast, moved at a steadier, lower-variance velocity (Supplementary Figure 1c), and its MSD scaled sub-diffusively (α = 0.85), consistent with movement constrained to the hyphal surface and punctuated by pauses and reversals (Figure 1d). Movement-direction histograms for both organisms were bimodally clustered at 0° and 180° relative to the hyphal axis (Supplementary Figure 1e), far from the uniform distribution expected under isotropic random motion, confirming that the axial alignment in Figure 1c reflect movement in both directions along the hyphal axis rather than a single dominant direction.

### Protist communities converge reproducibly to a single species while bacterial assembly remains largely stochastic

Alluvial diagrams constructed from Emu genus-level abundance tables revealed pronounced and directional community restructuring across successive hyphal-baiting generations in both protist (18S) and bacterial (16S) communities (Figures 2a, 2b). The rhizosphere protist community was compositionally complex, with diversity distributed across *Eocercomonas* (4.1%), *Flamella* (3.0%), *Cercomonas* (1.5%), and *Hartmannella* (1.3%), while *Paracolpoda* was detected at only 0.6% relative abundance. Following a single round of hyphal baiting, the community collapsed toward three dominant genera: *Acanthamoeba* (16.6%), *Paracolpoda* (13.5%), and *Paracercomonas* (6.9%), with marked reduction of rhizosphere-associated taxa. Through phylogenetic analysis, the dominant *Paracolpoda* consensus sequence was identified as belonging to a clade within the species *Paracolpoda steinii*, in a genus closely related to *Colpoda* (Supplementary figure 2a). The *Paracercomonas* consensus sequence formed a well-supported clade with *P. marina* (Supplementary Figure 2b).

By G2, *Paracolpoda* became the dominant protist genus, while *Acanthamoeba* and *Paracercomonas* declined to undetectable levels, indicating a continued selection imposed by prolonged hyphal association (Figure 2b; Supplementary Figure 3c). This endpoint was reproducible across independent replicates, with all five G2 microcosms converging on the same dominant genus (Supplementary Figure 3c). The bacterial (16S) community underwent a complementary transition from a *Bacillus*, *Domibacillus* and *Azotobacter* dominated rhizosphere toward Enterobacteriaceae dominance. *Enterobacter* increased from undetectable in the rhizosphere to 49.1% in G1 and 72.6% in G2. This progressive enrichment across successive hyphal transfers mirrors the deterministic recruitment dynamics previously observed in the FOV4 hyphosphere (Antony-Babu et al., 2025).

The assembly-process inference separated the two kingdoms. For bacteria, most comparisons fell within β-nearest taxon index (β-NTI) ≤ 2 and were classified by RC_bray as undominated (drift), including rhizosphere-G1 (100%), rhizosphere-G2 (84%; 16% dispersal limitation), G1-G2 (96%) and G2-G2 (89%; 11% dispersal limitation) (Figure 2c). The exception was within G1, where 56% of comparisons indicated homogeneous selection (β-NTI < -2), suggesting that the first round of hyphal baiting imposed a transient deterministic filter that gave way to drift-dominated assembly by G2. For protists, all estimable comparisons were classified as drift (Figure 2c), but these statistics were uninformative at the final generation, because observed richness had fallen to roughly two genera and every pairwise G2 comparison shared an identical taxon set, leaving β-NTI undefined (n = 0/10; Figure 2c). The reproducibility of the G2 endpoint across independent replicates nonetheless indicates consistent selection, which phylogenetic turnover statistics cannot detect when communities share an identical taxon set. Alpha diversity fell sharply from rhizosphere to G1 in both communities and remained low through G2 (Supplementary Figure 3a), and beta diversity ordination separated rhizosphere, G1, and G2 into three well-resolved clusters (PERMANOVA, bacteria pseudo-F = 542.3, R² = 0.99, p = 0.001; protist pseudo-F = 1030.8, R² = 0.99, p = 0.001; Supplementary Figure 3b), confirming that each hyphal-baiting generation occupied a distinct region of community composition space.

Linking genus identity to inferred ecological function (Figure 2d) showed that the G1 community spanned bacterial grazing, cyst formation and N & P mineralization, with smaller contributions from hyphal surface association, endosymbiont hosting and pathogen dispersal. By G2, functional representation was distributed roughly evenly across cyst formation, bacterial grazing, and N & P mineralization (each ∼23% of literature-weighted abundance), with smaller contributions from selective grazing (18%) and hyphal surface association (13%), because the G2 protist community is dominated by a single genus, this even distribution reflects the range of functions attributed to that genus in the literature rather than functional diversification across taxa.

### Topology alone is sufficient to guide *P. steinii* motility, independent of living fungal hyphae

To test whether the surface-guided motility observed in Figure 1 depends on any property specific to the living fungus, we tracked the ciliate *P. steinii* on both FOV4 hyphae and an inert glass rod (Figure 3a; Supplementary Videos 3-4). *P. steinii* was observed on both surfaces, but with markedly different kinematics, median cell speed was about 35 fold higher on the rod than on hyphae (Figure 3b). On the glass rod, individual trajectories were consistently biased along the fiber’s long axis (Figure 3c, left; Supplementary Videos 3), and mean angular deviation from that axis (36.3°) was significantly lower than the 45° expectation under random orientation (p = 0.003; Figure 3d, left). On FOV4 hyphae, trajectories were visibly more tortuous and branch-exploring (Figure 3c, right; Supplementary Videos 4) and angular deviation from the nearest of several local hyphal branches did not differ significantly from its condition-specific random expectation (17.2°; p = 0.213; Figure 3d, right). Path straightness was correspondingly low on both surfaces (rod, 0.31 ± 0.16; hyphae, 0.24 ± 0.34; p = 0.617; Supplementary Figure 4a), a pattern consistent with frequent branch exploration on hyphae rather than the simple rail-following seen on the geometrically uniform rod. Together, these results indicate that linear surface geometry alone is sufficient to bias the direction of *P. steinii* movement.

**Figure 3.**
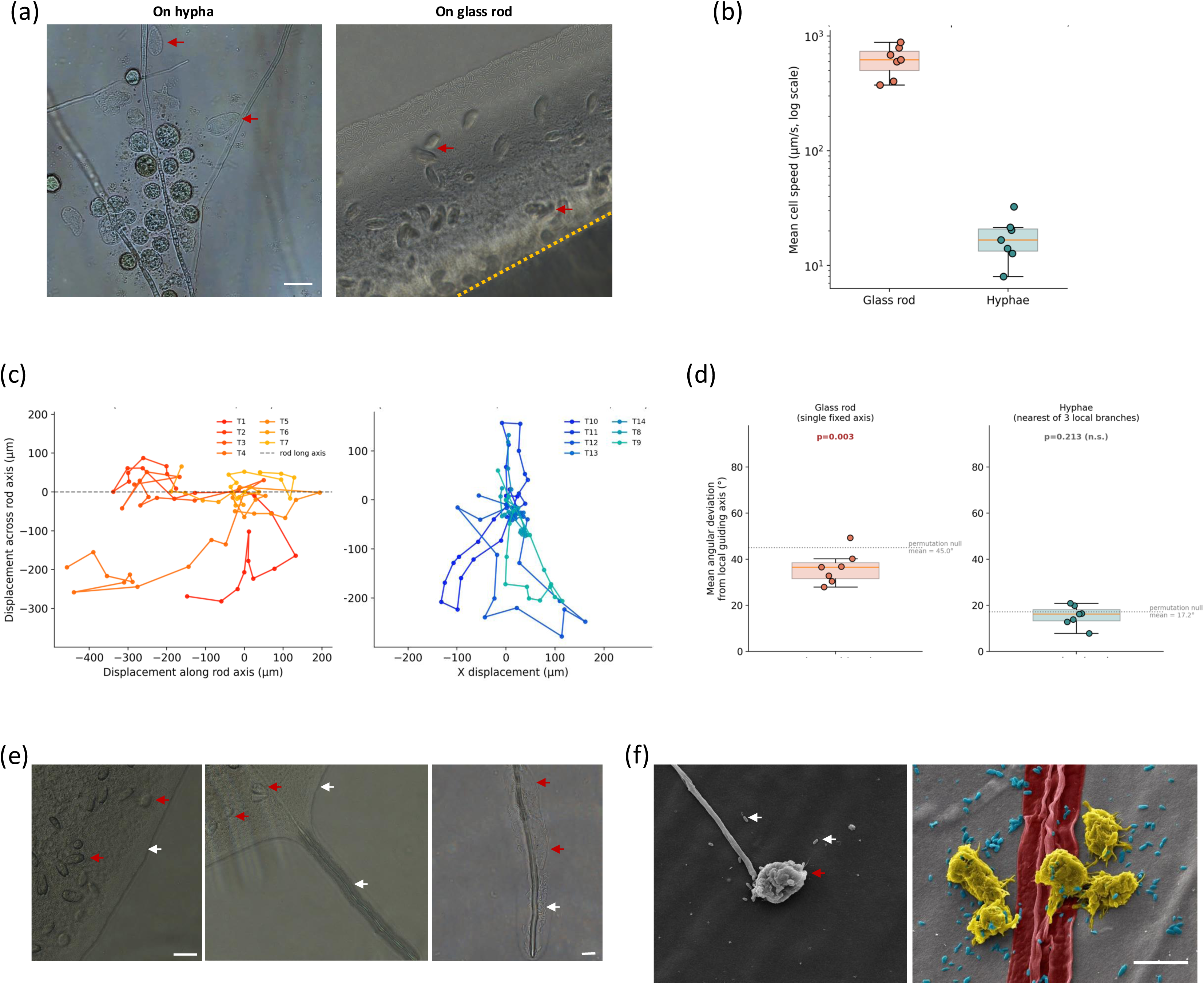
Topology-guided surface motility of *P. steinii* on fungal hyphae and an inert glass rod. **(a)** Representative micrographs of *P. steinii* (red arrows) on FOV4 hyphae (left) and on an inert glass rod (right; dashed yellow line - rod edge), demonstrating association with a fiber-like surface independent of fungal chemistry. Scale bar, 10 µm. **(b)** Mean cell speed of *P. steinii* on the glass rod versus on hyphae (n = 7), log scale. Horizontal bars, group medians. **(c)** Individual *P. steinii* trajectories on the glass rod (left, rotated to align the rod’s long axis horizontally, n = 7) and on hyphae (right, native orientation, translated to a common origin, n = 7). **(d)** Mean angular deviation of *P. steinii* movement: the rod’s single fixed long axis (left) or the nearest of three hyphal branches (right). Dashed lines indicate the random-expectation baseline for each geometry. Movement on the rod was significantly more aligned than random (permutation test, n = 7, p = 0.003); alignment on hyphae did not differ significantly from random (n = 7, p = 0.213). **(e)** Time-course micrographs illustrating community assembly on hyphae: bacteria (white arrows) and *P. steinii* (red arrows) co-occurring in the bulk colony (left); bacteria colonizing an extending hyphal strand ahead of *P. steinii* arrival (middle); *P. steinii* subsequently migrating onto colonized hyphae (right). Scale bars, 10 µm. **(f)** Scanning electron micrographs confirming direct physical association between bacteria and *P. steinii* with fungal hyphae. Left: A *P. steinii* cell (red arrow) at the tip of an isolated hyphal strand with bacteria around (white arrows). Right: Multiple *P. steinii* cells together with rod-shaped bacterial cells directly on a hyphal surface (Pseudo colors:. *Enterobacter* sp. - blue; *P. steinii -* yellow; FOV4 - red). Scale bar, 20 µm.

Independent of the kinematic assay, time-course microscopy illustrated a sequence of colonization, with bacteria and *P. steinii* initially co-occurring within the bulk colony, bacteria colonizing an extending hyphal strand in advance of protist arrival and *P. steinii* subsequently migrating onto the now bacteria-colonized hypha (Figure 3e). Scanning electron microscopy confirmed direct physical contact between *P. steinii*, bacteria and hyphal surfaces, including cells positioned at hyphal tips and multiple *P. steinii* cells co-located with rod-shaped bacterial cells directly on the hyphal surface (Figure 3f; Supplementary Figure 4b-e).

### A fungal bridge is required for cross-gap dispersal of both bacteria and protists and protist presence limits bacterial dispersal

To test whether fungal hyphae are necessary for multi-kingdom dispersal, we established paired donor-recipient microcosms separated by a 3 mm air gap under four donor treatments (see methods) and scored recipient-side colonization after 72 h (Figure 4). In treatments lacking FOV4 (protist + bacteria alone or bacteria alone), where no fungal hyphal bridge was formed, neither organism were detected in the recipient microcosm in any replicate (Figure 4d). Where FOV4 was present, hyphae reliably bridged the gap regardless of which other organisms were co-inoculated, including in a fungus-only control, confirming that hyphal bridging itself does not require co-inoculation with bacteria or protists (Figure 4b, c).

**Figure 4.**
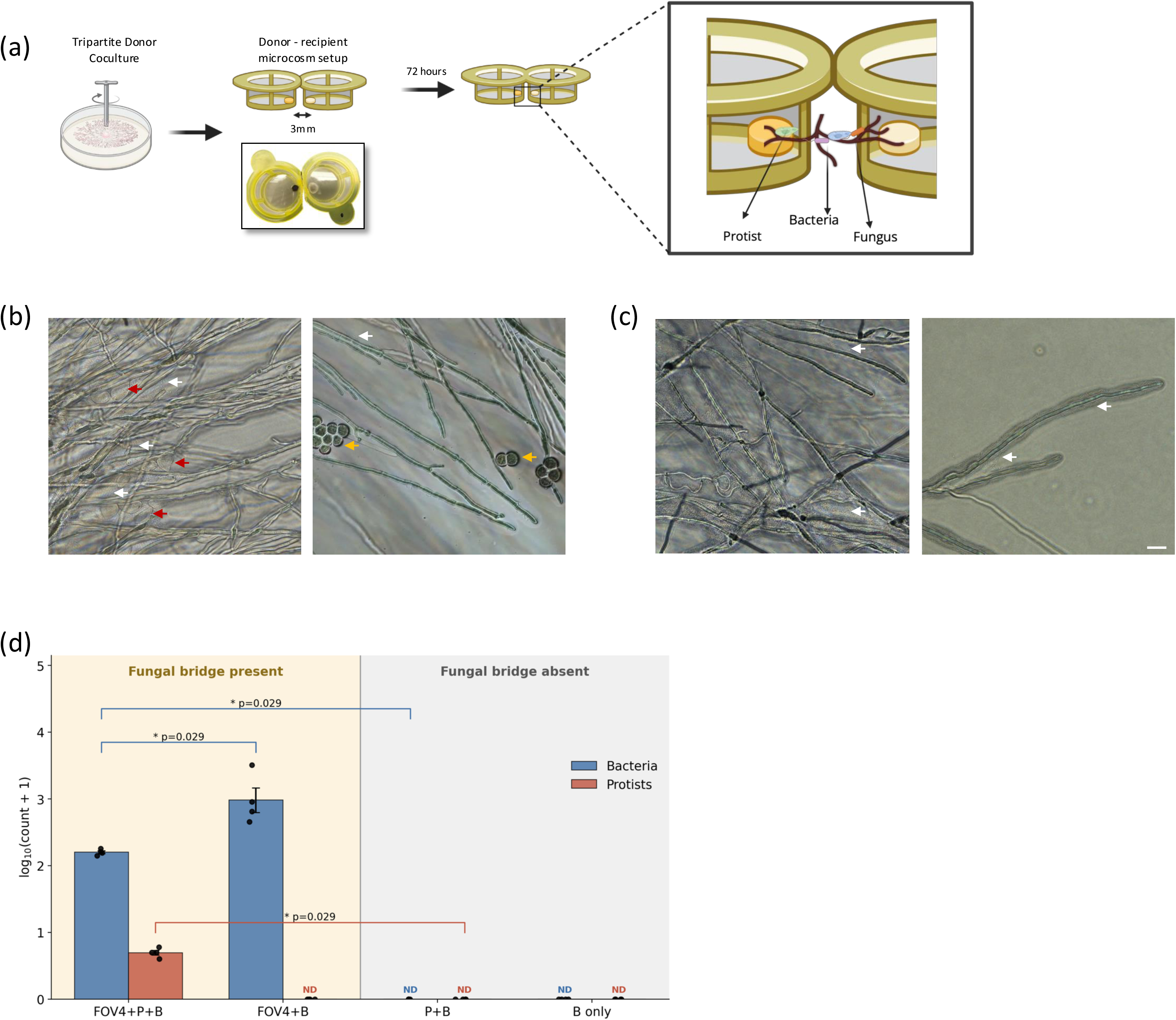
Fungal hyphae mediate multi-kingdom dispersal of bacteria and protists across a physical gap. **(a)** Schematic of the dual-microcosm dispersal assay. Donor co-cultures were prepared on 0.1x PDA for each treatment, and a 1 cm donor disc was placed in the donor strainer, facing a sterile 0.1x PDA disc in the recipient strainer across a 3 mm air gap. Coverslips at the base of each strainer prevented hyphae from growing underneath. After 72 h at 28 °C, the gap was inspected for hyphal bridging and cross-microcosm colonization by protists (green, blue) and bacteria (orange, purple), as illustrated in the inset. **(b)** FOV4 + protist + bacteria treatment. Bacterial (white arrows), protist (red arrows) cells and protist cysts (orange arrows) are visible at the hyphal entry point on the recipient agar plug, indicating active cross-gap colonization by both kingdoms. **(c)** FOV4 + bacteria(no protist). Bacterial cells (white arrows) are visible along the hyphal network within the recipient plug, confirming bacteria alone can exploit hyphae for spatial dispersal independent of protist presence. Scale bar, 10 µm. **(d)** Abundance of bacteria and protists recovered from the recipient microcosm across treatment groups, grouped by presence or absence of a fungal bridge. Bars show mean log₁₀(count + 1) (n = 4 per group, overlaid as points). ND, not detected. In the absence of a fungal bridge (P + B, B-only), neither organism was detected in the recipient microcosm. Among fungus-bridged treatments, recipient colonization in the FOV4 + protist + bacteria treatment was significantly greater than in the no-fungus control for both bacteria and protists (exact two-tailed Mann-Whitney U, p = 0.029 for each). Bacterial recovery was additionally 8.2-fold higher when protists were absent from the donor culture (FOV4 + bacteria vs. FOV4 + protist + bacteria, p = 0.029).

Both bacteria and protists were recovered from the recipient microcosm only when a fungal bridge was present (Figure 4b, c) and this dependence was statistically robust as recipient colonization in the FOV4 + protist + bacteria treatment was significantly greater than in the no-fungus control for both bacteria and protists (Mann-Whitney U, p = 0.029 for each; Figure 4d). Bacterial recovery was itself sensitive to protist presence as recipient bacterial counts were 8.2-fold higher when the donor culture contained bacteria and FOV4 alone than when the protist was also present (p = 0.029), even though hyphal bridging occurred in both treatments. This is consistent with protist grazing reducing bacterial abundance in transit or after arrival.

## Discussion

Fungal highways have been studied predominantly as routes for bacterial dispersal (Kohlmeier et al., 2005; Warmink et al., 2011; Jansa & Hodge, 2021). Hyphae are known to enhance protist colonization and dispersal range across soil pore space (Mafla-Endara et al., 2021), but that evidence is at population-level and it did not resolve or quantify trajectory-level movement directly along the hyphal surface itself, nor it was compared with hyphae or with an inert structure of similar geometry. We show that a soil ciliate, *Paracolpoda steinii,* exploits fungal hyphae as a locomotory surface, and that an inert substrate sharing the physical geometry of a fungal hypha is sufficient to guide this movement as *P. steinii* aligned significantly with the long axis of a chemically and biologically inert glass rod, producing a qualitatively similar pattern of axial alignment to that measured for bacteria and the flagellate protist *Paracercomonas marina* on real hyphae (Figure 1, Figure 3). Our results indicate that fibre-like geometry is sufficient for directional guidance even without fungal-specific substrate identity, extending the fungal highway concept beyond a single kingdom, although it does not exclude additional contributions from fungal surface chemistry or biological activity on living hyphae. Because *P. steinii* is the same species tracked in the trophic-filtering assay below, this geometry-sufficiency result also identifies the physical basis of the predator’s access to its prey’s dispersal route, rather than merely coinciding with it. Together with the trophic-filtering effect described below, this reframes the fungal highway not simply as a bacteria-specific transport route now shown to extend to a second kingdom, but as shared infrastructure whose traffic is itself reshaped by interactions among the kingdoms using it.

On the glass rod, *P. steinii* trajectories were consistently biased along a single, geometrically simple axis. On hyphae, movement relative to the nearest of several local branch angles was statistically indistinguishable from random, even though path straightness was equally low on both surfaces. These trajectories nevertheless remained confined to the hyphal surface despite the absence of a preferred local branch orientation. Rather than showing an absence of guidance, we interpret this as evidence that hyphal topology offers *P. steinii* a more complex physical environment consistent with evidence that pore and branch geometry alone can shape microbial growth and movement independent of chemistry (Arellano-Caicedo et al., 2021), and with a synthetic glass-fibre network similarly reproducing fungal-mediated bacterial dispersal and community shifts in a different microbiome (Zhang et al., 2018) one with real branch points and unlike the rod, a live microbial community so that the resulting tortuous trajectories may reflect exploratory behavior associated with foraging, though confirming this would require behavioral metrics beyond angular alignment.

Across two generations of hyphal baiting, the protist community collapsed from a diverse rhizosphere assemblage to three dominant genera; *Acanthamoeba*, an amoeba with well-documented bacterivorous and bacteria-hosting ecology (Khan & Siddiqui, 2014), *P. steinii*, and *Paracercomonas* after one baiting round and to *Paracolpoda* alone by G2 (Figure 2b). Colpodida species are among the most widespread and abundant soil ciliates, with a conserved life history that includes both feeding and cyst stages (Li et al., 2024) and have been shown to promote plant growth through bacterially mediated phosphate transport in other systems (Zhang et al., 2022) a function consistent with the N & P mineralization role inferred for this genus here (Figure 2d). β-NTI and RCbray analysis (Stegen et al., 2013, 2015), indicated that bacterial convergence was dominated by drift, with homogeneous selection detectable within G1 (Figure 2c). For protists these statistics were not computable at G2, because replicate communities had collapsed to an identical taxon set. This partitioning of stochastic and deterministic processes follows established practice in microbial community assembly (Nemergut et al., 2013; Dini-Andreote et al., 2015; Zhou & Ning, 2017). Inferred ecological functions shifted across same generations. The G1 was biased toward bacterial grazing, an established mode of top-down bacterial control by soil amoebae (Weekers et al., 1993) together with cyst formation and N & P mineralization. By G2, the representation was spread more evenly across grazing, mineralization and cyst formation (Figure 2d), consistent with the hyphosphere acting as a progressively narrowing filter that selects, across generations, for taxa favored within the hyphosphere.

A fungal bridge was required for either bacteria or protists to colonize a recipient microhabitat across a 3 mm air gap and without it, no colonization occurred in any replicate (Figure 4). But bacterial colonization of the recipient side was substantially lower when *P. steinii* was present in the donor culture than when it was absent, suggesting grazing by *P. steinii*, an established bacterivore (Geisen et al., 2016; Martins et al., 2022), reducing the number of bacteria that complete the crossing, though we did not directly observe grazing in this assay and our design cannot rule out other explanations, such as resource competition between the co-inoculated organisms. If grazing is the operative mechanism, it implies that fungal highways are not a passive conduit that simply reflects donor-community composition at the far end, but a filter whose output may be shaped by interactions among the organisms traveling on it. Protist community composition is itself a strong indicator and determinant of plant performance in other systems (Guo et al., 2021), and community composition and bacterial interactions are known to shape *Fusarium* disease outcomes more broadly, from suppressive-soil microbiomes to rhizosphere community shifts during infection (Bautista et al., 2023; Yang et al., 2023), underscoring that microbiome assembly, not the pathogen alone, governs disease-relevant outcomes in this genus. This bears directly on a pathogenic, rather than saprotrophic or mutualistic, fungal highway as FOV4 recruits a hyphosphere whose bacterial members can promote its growth in a strain-specific manner, with a possible but still unresolved role in virulence (Antony-Babu et al., 2025). If protist grazing constrains which bacteria persist there before or during host infection, protist activity could influence assembly of that pathogen-associated hyphosphere, a route not addressed by prior characterizations of this pathogen, raised here as a possibility rather than a demonstrated disease consequence. One limitation of here is that the ecological function assignments in Figure 2d are inferred from published evidence for the corresponding genera rather than measured directly in this study. They therefore represent putative functional potential rather than experimentally validated activities.

The finding that an inert glass rod reproduced directional guidance suggests that fungal highways can function as physical guidance structures, while biological interactions modify the ecological consequences of that guidance as the topology experiment supports the guidance itself. The dispersal assay shows that what travels along it, and how much, is then shaped by the organisms sharing that surface. Previous studies have largely interpreted fungal highways as an opportunity for bacterial dispersal. Our findings show that a eukaryotic cell, orders of magnitude larger than bacteria and reliant on entirely different locomotory machinery, can nonetheless use the same hyphal surface as a directional guide extending the fungal highway from a bacteria-specific phenomenon to one that spans kingdoms. Together with the finding that community assembly and dispersal along these highways are further shaped largely by drift and by interactions between protists and bacteria and this identifies pathogenic fungal highways as dynamic ecological corridors rather than passive, bacteria only routes of microbial transport. Extending this framework to other pathogenic and mutualistic fungal systems and to a wider range of protist locomotory strategies, will help clarify how general this coupling is across the fungal highways that structure soil microbial life.

## Conclusion

This study shows that fungal hyphae are not a bacteria-exclusive dispersal route, but a shared surface used directionally by organisms from different kingdoms. Protists, despite being far larger than bacteria and relying on entirely different locomotory machinery, moved along hyphae with strong axial alignment, and the geometry of a hypha alone was sufficient to bias protist movement. Community composition across successive hyphal-baiting generations changed reproducibly across generations, with bacterial assembly dominated by drift and protists converging on a single ciliate species reframing the hyphal surface as a site of community assembly, not just transit. Within a pathogenic system specifically, the protist’s presence cut bacterial arrival at a distant microhabitat roughly eightfold, a pattern matching predation rather than passive co-transport. Read together these results position the fungal highways of a plant pathogen as multi-kingdom corridors where physical guidance may shape which organisms travel and where interactions among those organisms along the way shape which communities form. The dual-microcosm design used here demonstrates that a fungal bridge is necessary, for cross-habitat colonization by either bacteria or protists, a distinction that offers a template for testing dispersal-dependence directly in other fungal systems, pathogenic or otherwise.

## Author Contributions

GA: conceptualization, validation, formal analysis, methodology, visualization, writing - original draft, review & editing. EDV: validation, formal analysis, writing - review & editing. SSP: formal analysis, writing - review & editing. ST: formal analysis, writing - review & editing. MB: writing - review & editing, resources, funding acquisition. LT: resources, writing - review & editing. SAB: conceptualization, validation, resources, funding acquisition, writing - review & editing.

## Acknowledgments

We thank the United States Department of Agriculture (USDA), College Station, Texas, for providing access to growth chamber facilities used in this study.

## Data Availability

All sequencing data generated in this study have been deposited in figshare under the accession https://doi.org/10.6084/m9.figshare.32896301

