## Supplementary data for "Hyphosphere Hunting Grounds: Fungal Hyphae Guide a Protist Predator to its Bacterial Prey"

Gayan Abeysinghe et al.

**This file includes:**

Supplementary Text
Figs. S1 to S4

**Other Supplementary Materials for this manuscript include the following:**
Movies Supplementary movie 1 to 4

SUPPLEMENTARY FIGURE LEGENDS:

**Supplementary Figure 1. Kinematic analysis of flagellate protist (left) and bacteria moving along FOV4 hypha (right).**

**(a)** Origin-centred trajectories of individual flagellate protist ($n = 7$, left) and bacterial ($n = 14$, right) tracks, translated to a common origin (filled circle). Terminal positions shown as filled points.

**(b)** Cumulative path length (µm) over time (s) for individual tracks. Tracks extend to 60s.

**(c)** Instantaneous velocity (µm $s^{-1}$) over time, smoothed with a Gaussian filter ($\sigma = 0.6$).

**(d)** Mean squared displacement (MSD, µ$m^2$) as a function of lag time $\tau$ (s) for individual tracks (coloured lines) and population-level power law fit (dashed black, MSD ∝ τ^α). Dotted grey line indicates Brownian diffusion ($\alpha = 1$). Flagellate protists showed subdiffusive motion ($\alpha = 0.85$), consistent with surface-constrained gliding; bacteria showed superdiffusive motion ($\alpha = 1.58$), consistent with persistent directed runs.

**(e)** Rose diagrams of movement direction relative to the hyphal axis (0° = tip-ward, 180° = base-ward). Dashed circle, expected frequency under random isotropic movement. Bimodal clustering at 0° and 180° confirms bidirectional surface-guided motility in both organisms.

**Supplementary Figure 2. Maximum Likelihood tree of dominant *Colpodida* and *Paracercomonas***

(a) Colpodida ML tree : Maximum likelihood (ML) phylogeny of the Colpodida dataset. ML bootstrap values > 75 and Bayesian posterior probabilities > 0.90 are shown above corresponding nodes. * indicates that ML bootstrap values are < 75. The consensus sequence for Colpodida X is shown in bold blue font.

(b) Paracercomonas ML tree: Maximum likelihood (ML) phylogeny of the Paracercomonadidae dataset. ML bootstrap values > 75 and Bayesian posterior probabilities > 0.90 are shown above corresponding nodes. * indicates that ML bootstrap values are < 75. The consensus sequence for *Paracercomonas marina* is shown in bold purple font.

**Supplementary Figure 3. Alpha diversity, beta diversity, and community composition across rhizosphere and hyphosphere generations.**

**(a)** Observed genus richness, Shannon diversity (H′), Simpson's index (1−D), and Pielou's evenness (J′) for the bacterial (16S, top) and protist/eukaryote (18S, bottom; after removal of plastid, mitochondrial, and host-plant *Gossypium* reads) communities across rhizosphere and hyphal-baiting generations. n = 5 per group. Kruskal-Wallis p-value shown above each panel; brackets denote pairwise Dunn's post-hoc comparisons significant after FDR correction (*p < 0.05, **p < 0.01, ***p < 0.001).

**(b)** Principal coordinates analysis (Bray-Curtis dissimilarity, genus level) of bacterial (top) and protist/eukaryote (bottom; plastid/mitochondrial/host reads removed) community structure across groups. Ellipses show 1.5 s.d. around group centroids. n = 5 per group. PERMANOVA (999 permutations) statistics reported in each panel title.

**(c)** Relative abundance of the top 12 bacterial (top) and protist/eukaryote (bottom) genera across rhizosphere and baiting-generation samples; remaining taxa grouped as "Other." n = 5 per group, labeled sequentially (R1-R5 / G1(1-5) / G2(1-5)).

**Supplementary Figure 4. Path straightness comparison and additional ultrastructural evidence of ciliate-hyphae association.**

**(a)** Path straightness (net displacement / total path length) of *P. steinii* on the glass rod versus hyphae (n = 7 tracks each). Horizontal bars, group means. No significant difference was observed between conditions (Mann-Whitney U, p = 0.617).

**(b-e)** Scanning electron micrographs of *P. steinii -Enterobacter*-hyphae association, complementing Figure 3f. (b) *P. steinii* cells at a hyphal branch point. (c,d) Lower-magnification fields showing multiple *P. steinii* hyphae contact points distributed across the hyphal network. (e) A second, independent example of *P. steinii* cells at a hyphal branch point.

SUPPLEMENTARY VIDEO LEGENDS:

**Supplementary Video 1. *Paracercomonas marina* movement along FOV4 hyphae**

**Supplementary Video 2. Bacterial movement along FOV4 hyphae**

**Supplementary Video 3. *Paracolpoda steinii* movement along an inert glass rod**

**Supplementary Video 4. *Paracolpoda steinii* movement on FOV4 hyphae**
