## Supplementary figures for "Hyphosphere Hunting Grounds: Fungal Hyphae Guide a Protist Predator to its Bacterial Prey"

### Supplementary Fig 1.

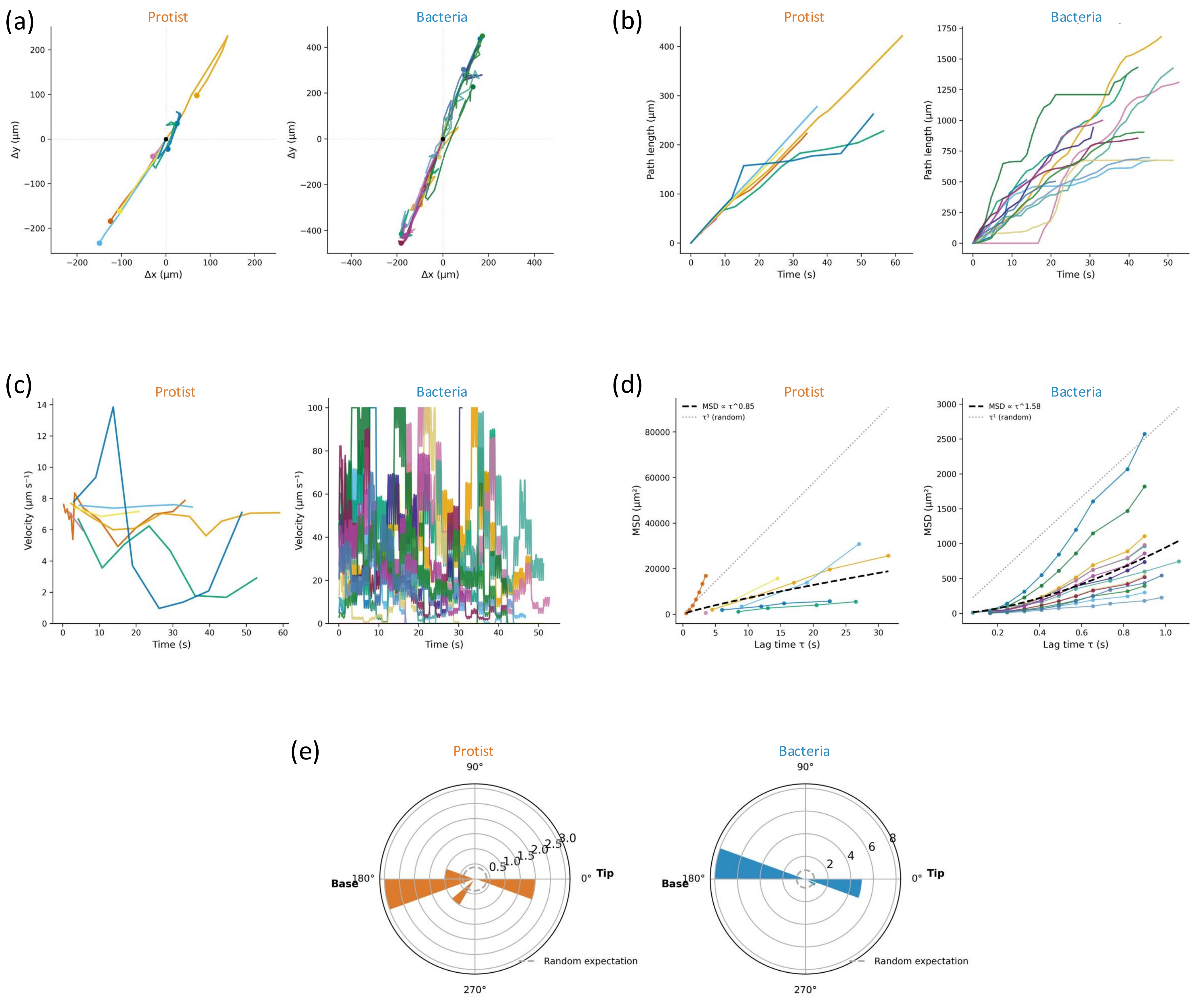

Supplementary Fig 2.

(a)

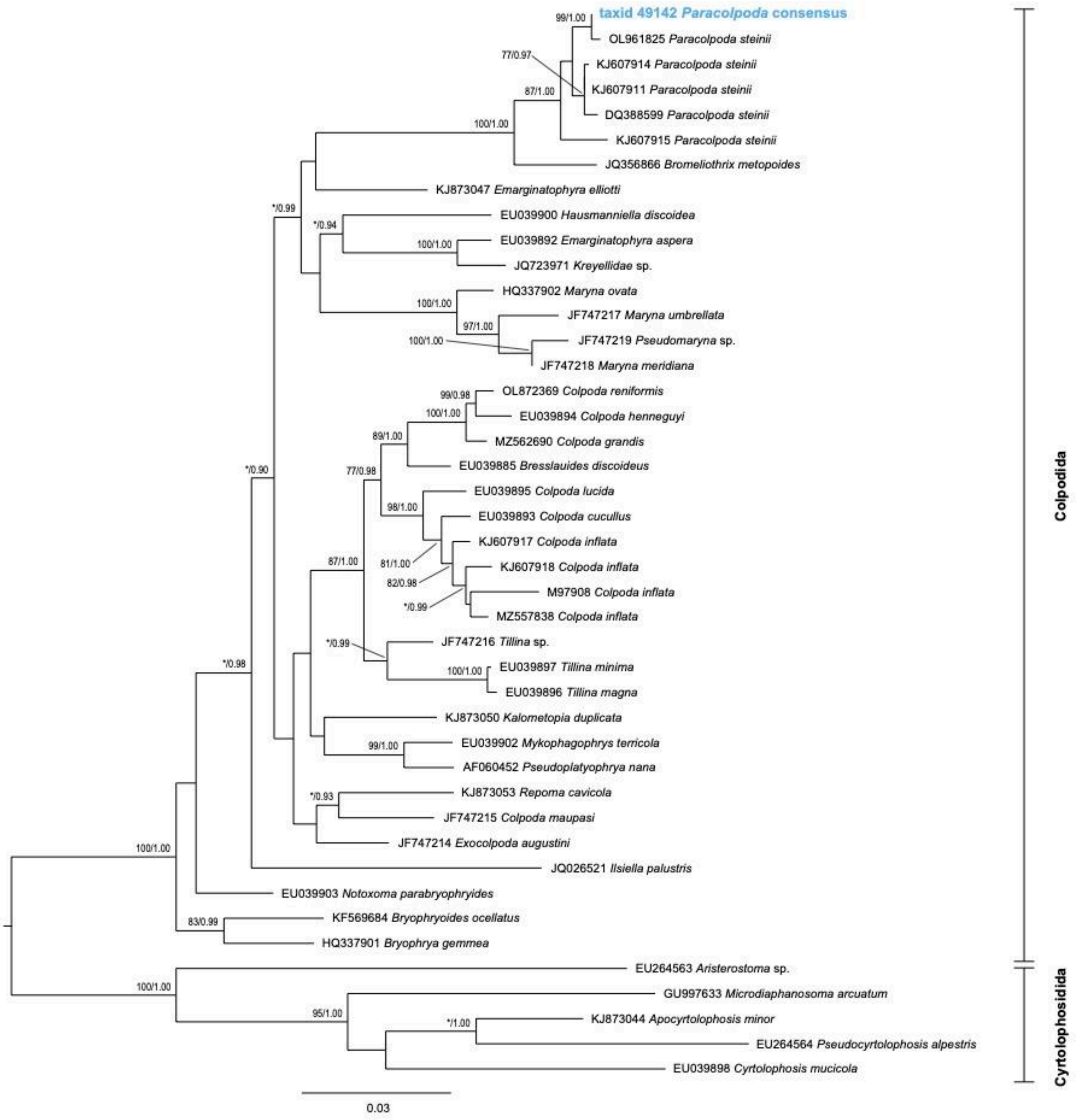

Supplementary Fig 2.

(b)

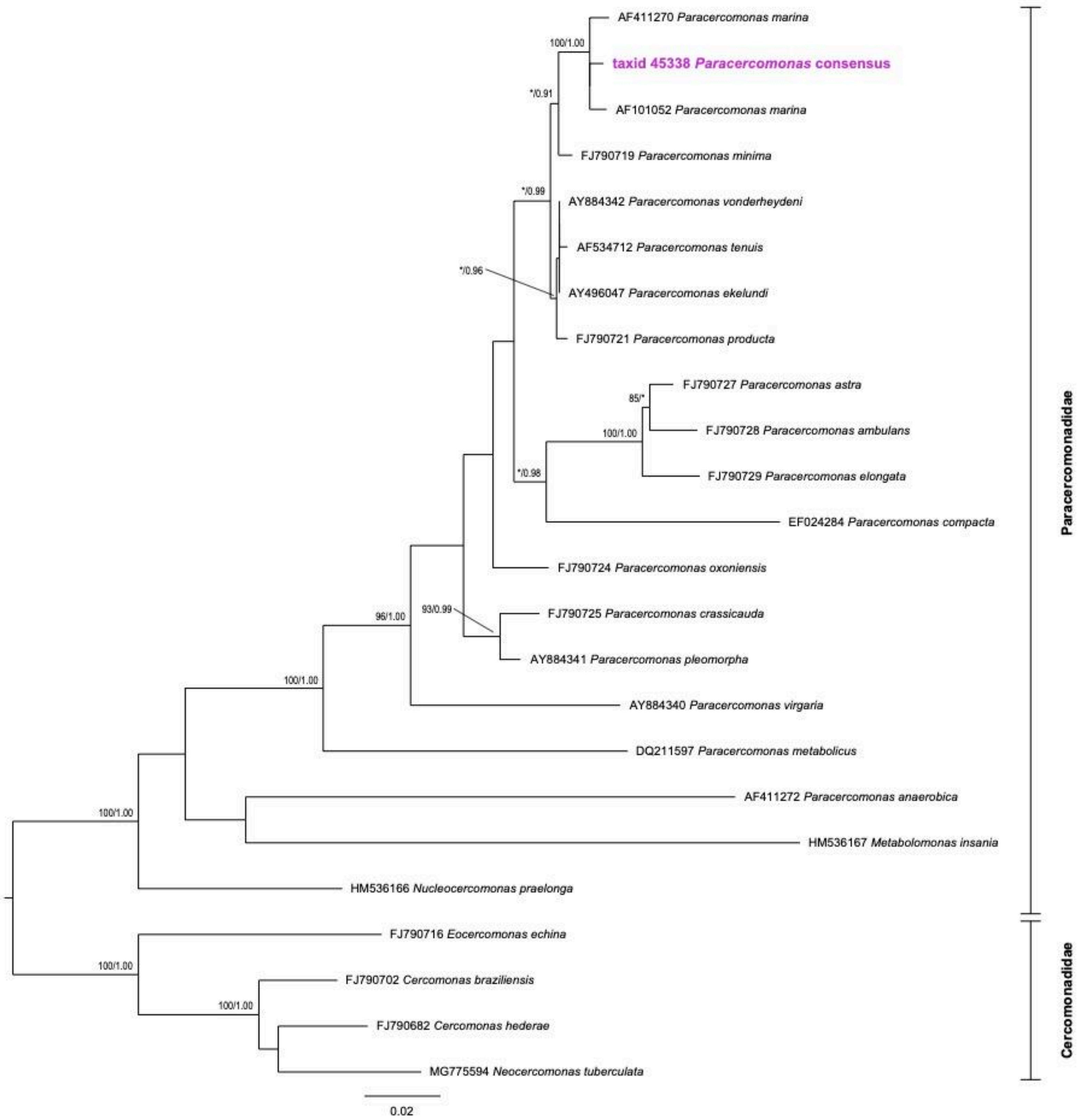

Supplementary Fig 3.

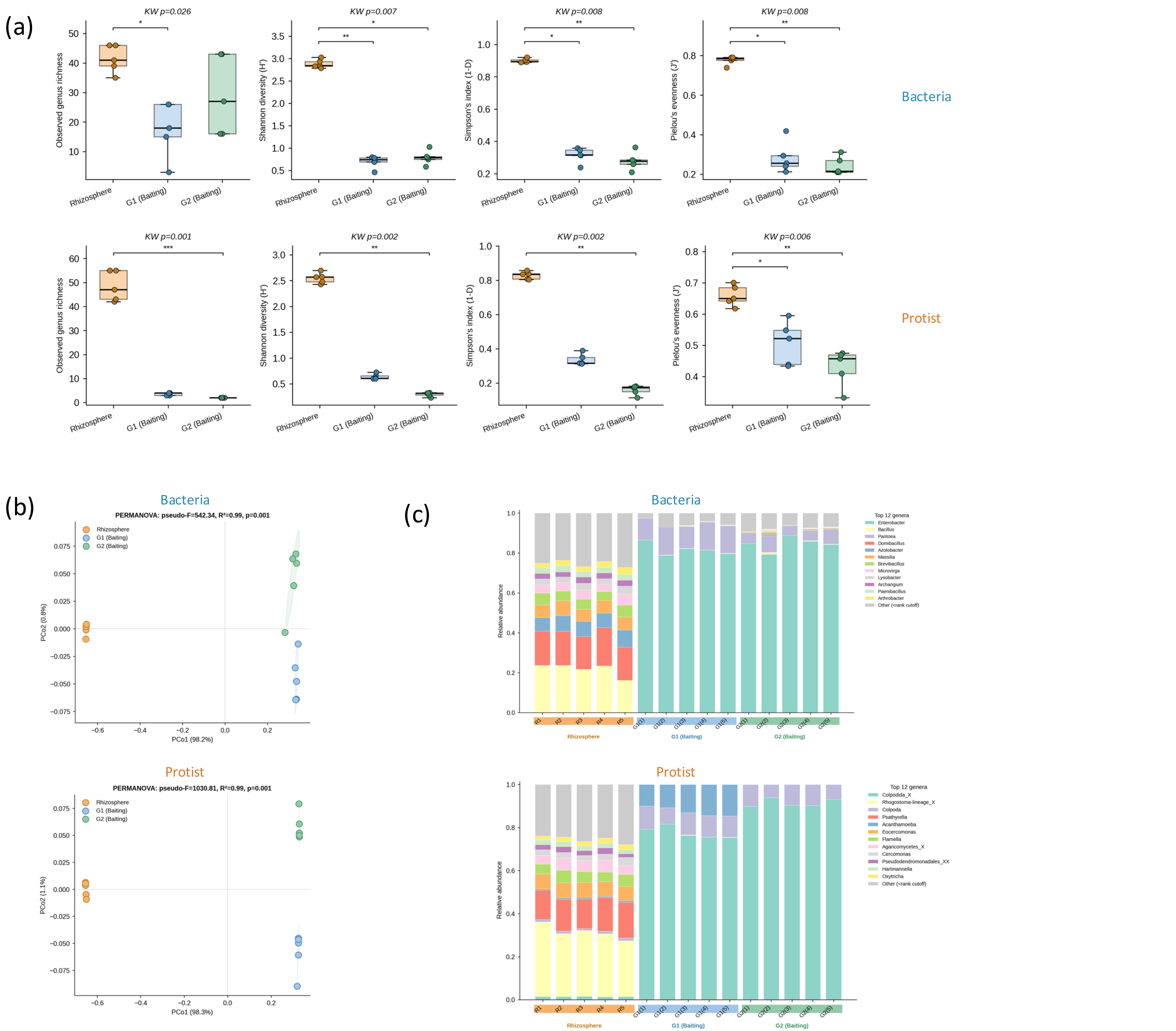

Supplementary Fig 4.

(a)

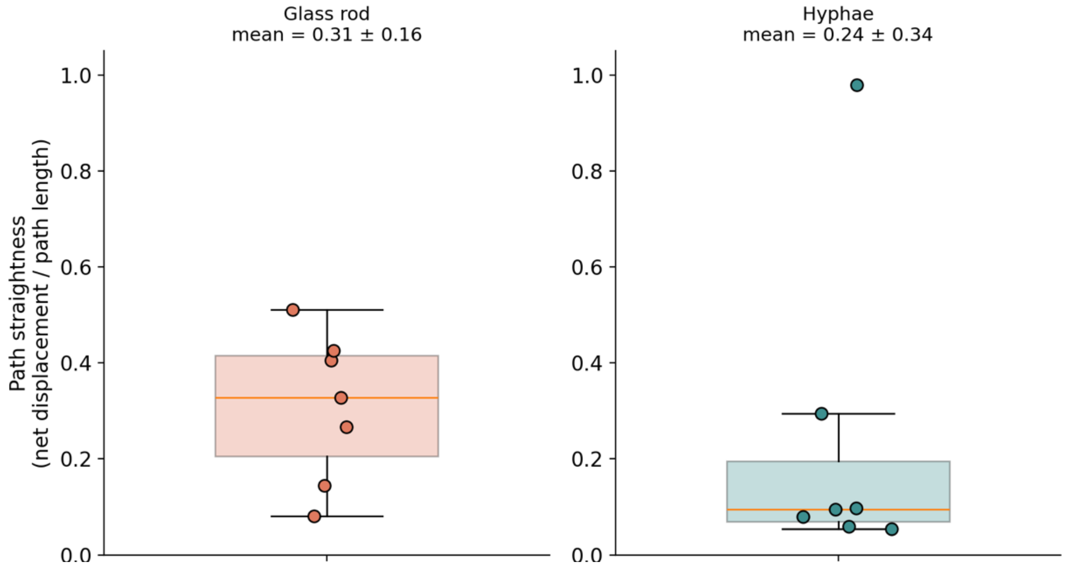


(b)

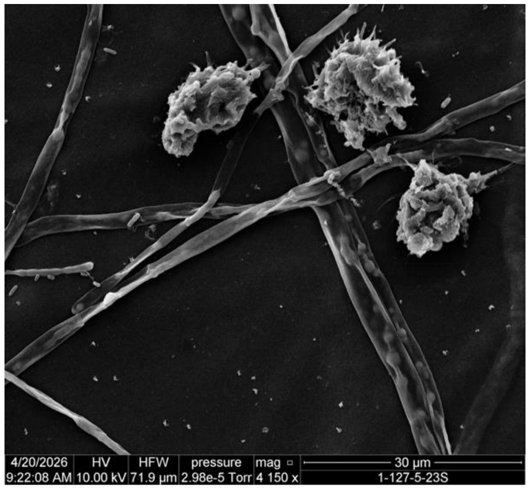


(c)

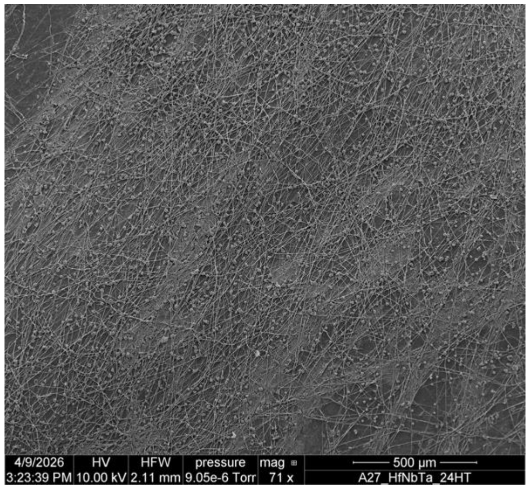


(d)

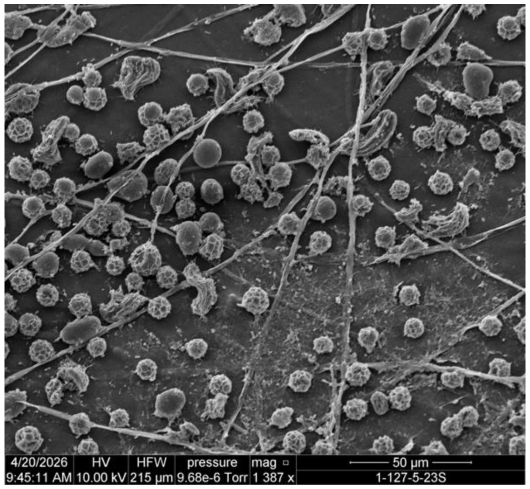


(e)

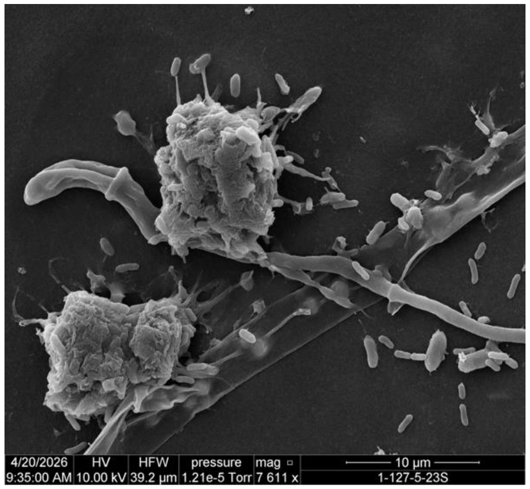
